# Piscis: a novel loss estimator of the F1 score enables accurate spot detection in fluorescence microscopy images via deep learning

**DOI:** 10.1101/2024.01.31.578123

**Authors:** Zijian Niu, Aoife O’Farrell, Jingxin Li, Sam Reffsin, Naveen Jain, Ian Dardani, Yogesh Goyal, Arjun Raj

## Abstract

Single-molecule RNA fluorescence *in situ* hybridization (RNA FISH)-based spatial transcriptomics methods have enabled the accurate quantification of gene expression at single-cell resolution by visualizing transcripts as diffraction-limited spots. While these methods generally scale to large samples, image analysis remains challenging, often requiring manual parameter tuning. We present Piscis, a fully automatic deep learning algorithm for spot detection trained using a novel loss function, the SmoothF1 loss, that approximates the F1 score to directly penalize false positives and false negatives but remains differentiable and hence usable for training by deep learning approaches. Piscis was trained and tested on a diverse dataset composed of 358 manually annotated experimental RNA FISH images representing multiple cell types and 240 additional synthetic images. Piscis outperforms other state-of-the-art spot detection methods, enabling accurate, high-throughput analysis of RNA FISH-derived imaging data without the need for manual parameter tuning.

## 1 Main

Modern fluorescence imaging techniques have enabled the study of complex biological systems at higher resolution and with more precision than ever before, and with it the demand for analytical tools that can perform at scale has grown. One specific area of interest is spot detection, owing to the development of numerous methods for single-molecule visualization (DNA^1^, RNA^2^, protein^3^) in which the targets are fluorescently labeled and detected as diffraction-limited spots. Spot detection allows for precise quantification of the abundance and localization of these molecules in single cells. In this rapidly evolving field, high-throughput spatial transcriptomics technologies^4–6^ such as MERFISH^7^ and SeqFISH+^8^ have emerged as powerful tools that can simultaneously measure the transcript abundance of large numbers of genes. These methods utilize combinatorial labeling and sequential imaging to capture and decode thousands of unique mRNA species *in situ*, generating vast datasets containing thousands to millions of cells, each with hundreds to thousands of fluorescent spots. Manual analysis is intractable, requiring an automatic and scalable workflow.

Various computational methods have been developed to automate spot analysis. These generally rely on bandpass filters via mathematical operators such as Laplacian of Gaussian^10^ or Difference of Gaussian^11^, coupled with a manual intensity threshold to differentiate high-frequency signals (spots) from low-frequency noise (background fluorescence). However, these methods often end up requiring *ad hoc* parameter tuning, which is time-consuming and may yield inconsistent results due to the subjective nature of manual adjustments. For instance, in two prominent software implementations, TrackMate (LoG)^12^ and RS-FISH (DoG)^13^, spot detection accuracy is highly dependent on the selected parameter combination (Supplementary Fig. 1), resulting in the need for extensive manual fine-tuning across different parts of the same dataset, such as different fields of view or even within the same field of view^2^. Moreover, when applied to images with high background noise, such as those with pronounced autofluorescent regions, TrackMate and RS-FISH are prone to detecting high numbers of false positives in these areas, severely distorting the final per-cell transcript counts (Fig. 1c-d).

**Figure 1:**
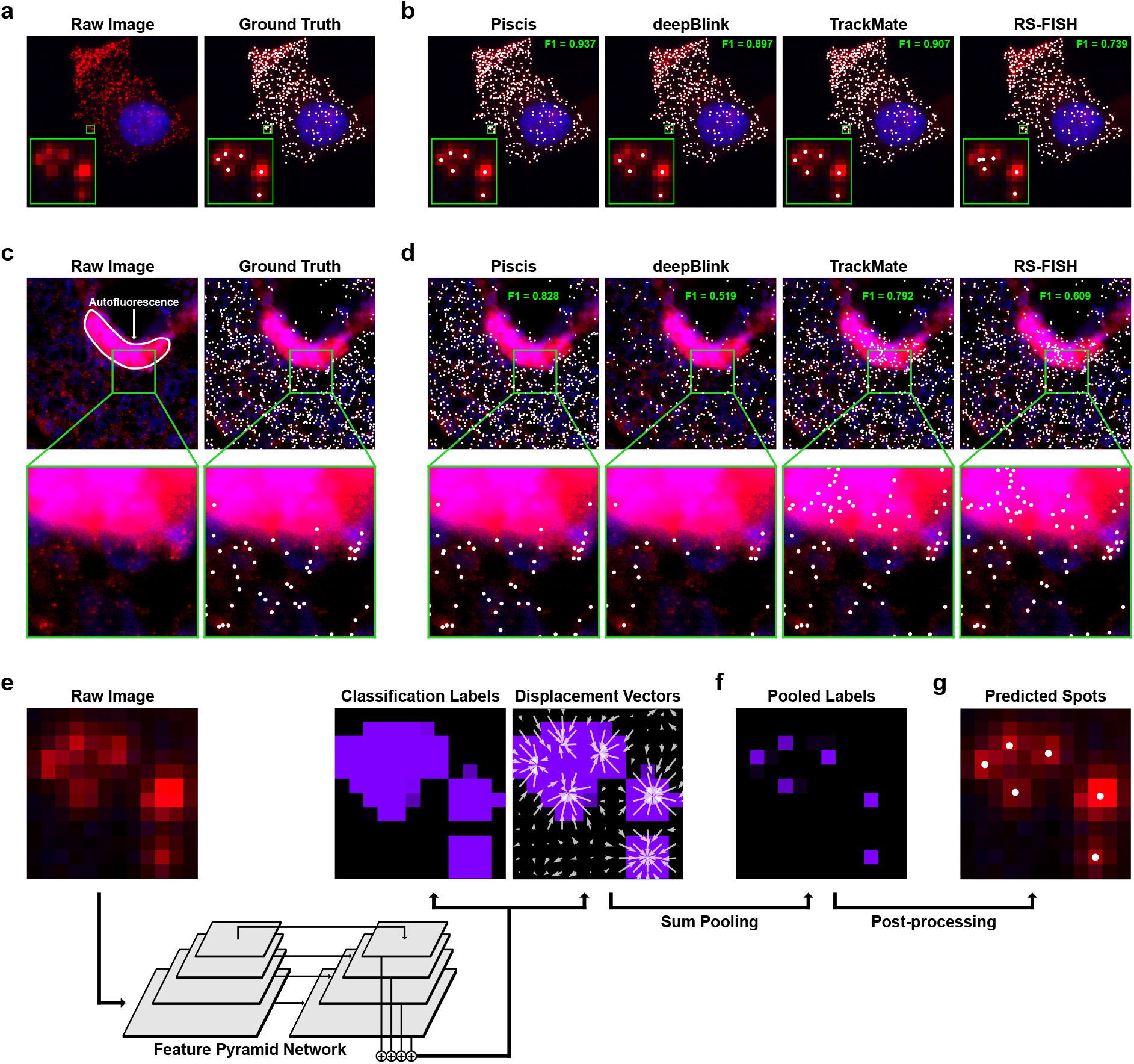
Piscis is a deep learning algorithm for accurate spot detection and localization. **a**, A single-molecule RNA FISH image (single z-level) of human fibroblast cells grown *in vitro* with DAPI-stained nuclei (blue), spots of single mRNA molecules for the gene *UBC* (red), and the corresponding manual ground truth annotations (white). **b**, Comparison between Piscis and other spot detection algorithms applied to the image from **a**. The output of each algorithm is labeled with its F1 score, a performance metric ranging from 0 to 1, with 1 representing perfect agreement with the ground truth. **c**, A clampFISH 2.0^9^ image (single z-level) of human melanoma cells grown in mice with DAPI-stained nuclei (blue), spots of single mRNA molecules for the gene *NGFR* (red), and the corresponding manual ground truth annotations (white). **d**, Comparison between Piscis and other spot detection algorithms applied to the image from **c**. The output of each algorithm is again labeled with its F1 score. **e**, Piscis uses a Feature Pyramid Network to predict classification labels and displacement vectors from a raw image. **f**, A deformable sum pooling operation is applied to generate more sharply peaked pooled labels. **g**, A post-processing step involving local maxima analysis and application of displacement vectors is performed to obtain predicted spots.

Recent advancements in deep learning have profoundly transformed many areas of biological image analysis, with neural networks achieving near-human-level performance in complex tasks like cell segmentation^14–17^ and tumor classification^18,19^. Such approaches thus hold considerable promise for automatic spot detection, with deepBlink^20^ and, more recently, Polaris^21^ being the most sophisticated implementations to date. However, a major challenge in training neural networks for this application is class imbalance: most image pixels represent the background, while relatively few pixels correspond to spots. This imbalance can bias the neural network toward the majority background (negative) class when using traditional training techniques, resulting in a high false negative rate for the underrepresented yet crucial spot (positive) class.

In principle, one method for training neural networks in the presence of class imbalance is to directly optimize the F1 score, which is a metric that balances the number of false positives and false negatives into a single value. However, the F1 score is generally non-differentiable, making it incompatible with optimization algorithms used to train neural networks. Others have implemented several alternative approaches, including variants of the cross entropy and Dice loss functions. Cross entropy loss is an information-theoretic metric that treats predicted and ground truth values as two probability distributions and quantifies their difference. Despite being a common choice for classification tasks, cross entropy loss is severely impacted by class imbalance. It is often computed pixel-wise and subsequently averaged or summed over the whole image, meaning that the negative class, with substantially more pixels than the positive class, has a larger influence on the loss value and is thus focused on more during training. This bias generally yields models with high classification accuracy for the negative class but low classification accuracy for the positive class, ultimately resulting in poor spot detection.

In contrast, Dice loss uses the Dice similarity coefficient, which measures the amount of overlap in the positive class between the predicted and ground truth values, normalized by the average number of positive instances from each. In this way, Dice loss does not rely on the absolute number of pixels from each class, making it less susceptible to class imbalance. Empirical comparisons have indeed shown that Dice loss and its variants are more effective than cross entropy loss and its variants, such as weighted cross entropy loss and focal loss, at handling class imbalance in numerous medical image segmentation tasks^22,23^. For this reason, deepBlink employed the Dice loss in its training process, but it was still insufficient for preventing underdetection in images with higher background noise and less well-defined spots, such as those from tissue samples (Fig. 1d).

We developed a deep learning algorithm called Piscis to address the challenge of spot detection. Piscis solves the class imbalance problem by training models using a loss function that we call the SmoothF1 loss, a differentiable approximation of the F1 score that can be optimized via gradient descent. When trained using the SmoothF1 loss, Piscis significantly outperforms other state-of-the-art spot detection methods without the need for manual parameter tuning. We anticipate that our approach for approximating the F1 score will have broad application to other problems exhibiting class imbalance.

## 2 Results

In order to achieve accurate spot detection and localization using deep learning, we first needed to encode spots in a representation that a neural network can learn from and predict. Following a similar strategy to that of deepBlink and Polaris, we used an auxiliary representation where the task is divided into predicting two types of images known as feature maps: 1. binary classification labels of each pixel as either spot or background and 2. displacement vectors that point each pixel to the nearest true spot center (Fig. 1e). The displacement vectors allow for the separation of nearby spots from each other because pixels in between spots will point either one way or the other. From the ground truth spots, we can generate the ground truth classification by labeling background pixels as 0 and spot pixels as 1. The number of pixels considered part of each spot depends on the loss function we use for model training. By default, our method Piscis uses a 3 × 3 region around the center of each spot. We can generate the ground truth displacement vectors by identifying the spot nearest to each pixel and then calculating the difference between their coordinates. If we could train a model to accurately predict these feature maps, we can subsequently process them to recover spot coordinates with subpixel localization accuracy, provided each pixel contains no more than a single spot. Note that even if a pixel contains two or more overlapping spots, they would be impossible to distinguish in practice due to the diffraction limit.

We trained a deep neural network to predict both the binary classification and displacement vectors from our raw images (Fig. 1e, Supplementary Fig. 2). The neural network architecture was a Feature Pyramid Network^24^ with an EfficientNetV2^25^ backbone, chosen for its previously demonstrated accuracy in image classification tasks and its superior parameter efficiency compared to other similar-performing models. This choice in network architecture was similar to those used in deepBlink, Polaris, and cell segmentation algorithms like Cellpose^15^. The neural network ultimately outputs three feature maps corresponding to the classification labels and the horizontal and vertical components of the displacement vectors. The classification labels predicted by the neural network alone can identify the image pixels that correspond to spots, but the identification of these regions alone is generally insufficient for discriminating individual spot centers as regions corresponding to nearby spots may merge together. We thus used the displacement vectors to distinguish the individual spots occupying these regions. We designed a deformable sum pooling operation, which shifts the classification label value of each pixel by its displacement vector to a new location (Fig. 1f). The resulting feature map, which we call the pooled labels, is more sharply peaked at spot centers, and height of each peak can be interpreted as a spot detection confidence score. After this sum pooling operation, we post-processed the final feature maps to obtain spot coordinates by performing local maxima analysis with a global threshold and further refining to subpixel localization accuracy via the application of predicted displacement vectors (Fig. 1g).

For model training and testing, we collected new or used existing experimental single-molecule RNA FISH images (*n* = 358; 23,222 spots) from a variety of cell types, including human fibroblasts, melanoma, lung adenocarcinoma, and macrophages (Fig. 2a). In our samples, multiple genes were labeled using probes coupled to various fluorophores and imaged across different fluorescent channels using either standard single-molecule RNA FISH or Hybridization Chain Reaction (HCR) RNA FISH. The objective of curating such a diverse dataset, representing a wide range of experimental and imaging conditions, was to train a model that can robustly generalize effectively to new, unseen images without extensive retraining. We further supplemented our dataset with synthetic images (*n* = 240; 24,768 spots) from the datasets used to train deepBlink. These datasets, named “Particle,” “Microtubule,” “Receptor,” and “Vesicle,” contained small particle-like objects with slightly different shapes and sizes compared to the spots in our experimental datasets. We included these datasets with the aim of further enhancing the robustness of our model.

**Figure 2:**
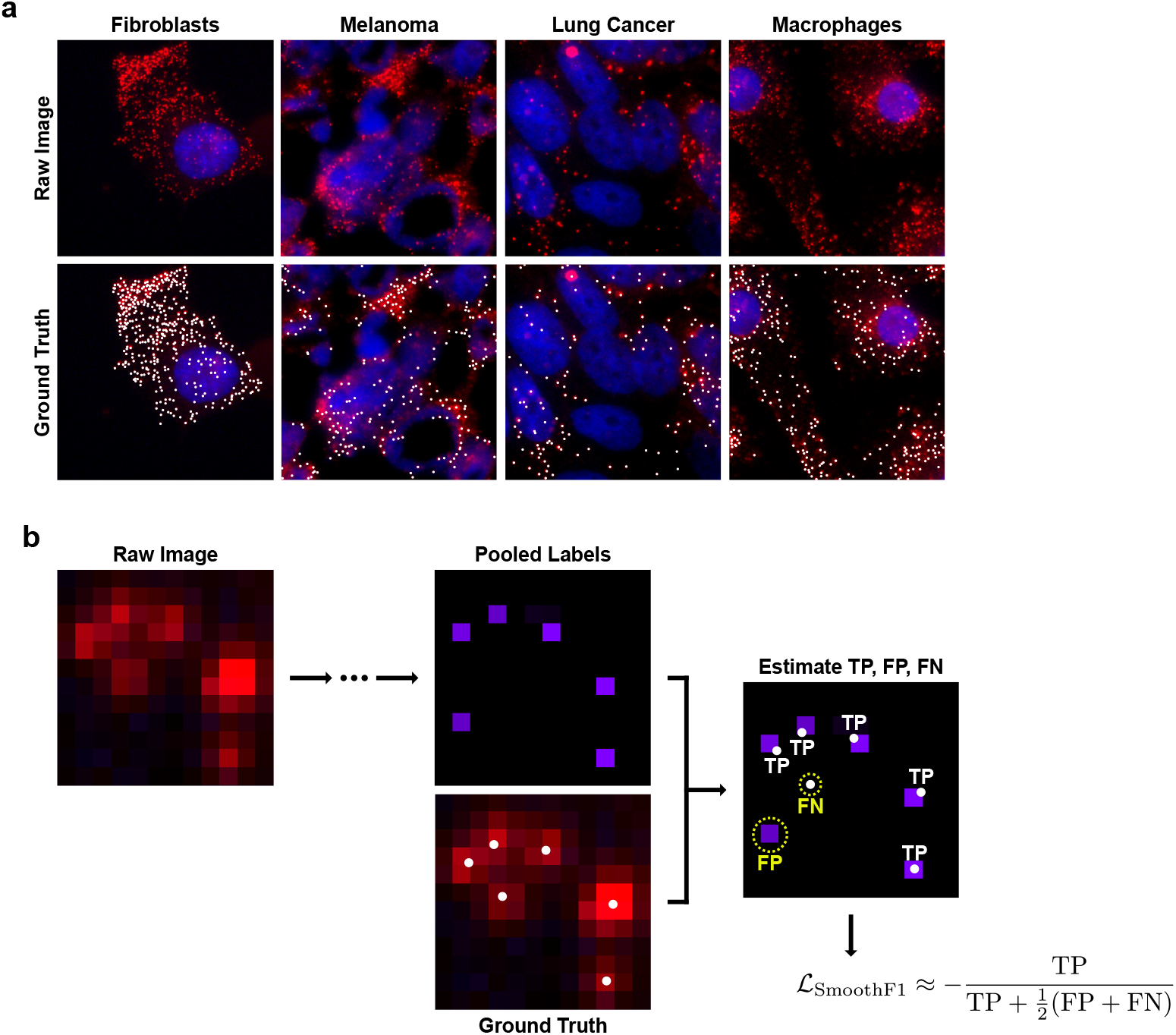
Piscis is trained on a diverse dataset using the SmoothF1 loss function. **a**, The dataset used to train and test Piscis included experimental RNA FISH images of human inducible fibroblast-like (hiF-T) cells^26,27^, WM989 human melanoma cells^28,29^, Calu-3 human lung adenocarcinoma cells^30^, and primary human monocyte-derived macrophages (hMDMs). Images are shown with DAPI-stained nuclei (blue), spots of single mRNA molecules (red), and the corresponding manual ground truth annotations (white). **b**, The SmoothF1 loss function enables direct optimization of the F1 score via an approximation of the true F1 score by assigning pixel values of the predicted pooled labels as either true positive (TP), false positive (FP), or false negative (FN) according to the manual ground truth annotations. Note that the pooled labels image shown here is a hypothetical example.

We manually annotated each spot in the experimental RNA FISH images using the custom software NimbusImage, an open-source, web-based platform for biological image analysis (see Methods). The coordinates of these manual annotations were refined by Gaussian fitting to obtain pseudo-ground truth for their subpixel localization. Synthetic images from the deepBlink datasets were pre-annotated, so we used their coordinates directly without additional processing. The final combined dataset of 608 images was then randomly split into training (*n* = 418; 33,821 spots), validation (*n* = 89; 6,977 spots), and testing (*n* = 91; 7,192 spots) sets.

To train Piscis, we needed to develop a custom loss function that addresses the class imbalance problem inherent to spot detection. In the specific case of spot detection, the majority of pixels are background and thus can dominate the sparser spot pixels when using standard loss functions. Although the Dice loss used in deepBlink has been previously shown to mitigate the effects of class imbalance and improve model performance, our testing still revealed a tendency for deepBlink to underdetect spots, leading to a high false negative rate (Fig. 1d).

In addition to the Dice loss, deepBlink artificially reduced class imbalance by grouping pixels into larger regions known as grid cells. Spots were then predicted on each grid cell rather than individual pixels. Larger grid cells thus reduced class imbalance but at the cost of lower detection accuracy in regions of high spot density, as the model cannot distinguish multiple spots within the same grid cell (Supplementary Fig. 3a-d). Among the 220 experimental images in our dataset containing at least ten spots, deepBlink’s default 4 × 4 grid cells would inadvertently omit more than 5% of the ground truth spots in 58 of these images during training (Supplementary Fig. 3e). These issues motivated us to develop a loss function that addresses class imbalance without relying on the pixel grouping inherent to grid cells.

Notably, one metric that does not suffer from class imbalance problems is the F1 score itself. The F1 score, defined as 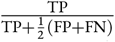, neatly incorporates true positives (TP), false positives (FP), and false negatives (FP) and indeed is often the metric used to measure overall model performance. However, the F1 score is typically a discrete function of the model predictions and hence is non-differentiable, making it impossible to use for model training as a loss function because gradient descent inherently requires differentiation. In the case of spot detection, the F1 score can only be calculated after several post-processing steps, including local maxima analysis to identify spot peaks and a combinatorial optimization algorithm to assign predicted spots to ground truth spots, both of which are non-differentiable. To address this issue, we developed the SmoothF1 loss function, a differentiable approximation of the F1 score for model training (see Methods). This approximation computes sums of the values in the pooled labels near, distant from, and missing at ground truth spots as proxies for the counts of true positives, false positives, and false negatives, respectively (Fig. 2b). We then used these values to compute an approximate F1 score. Given that the F1 score increases with improving model performance while loss functions decrease by convention, we thus define the SmoothF1 loss as the negative of this approximated F1 score.

To evaluate the effectiveness of the SmoothF1 loss function, we trained Piscis on our combined dataset and compared its performance on the testing set using the F1 score against deepBlink, TrackMate, and RS-FISH (Fig. 3a). To ensure a fair comparison, we retrained deepBlink on our combined dataset using multiple grid cell sizes and chose to represent the model with the optimal size of 2 × 2 pixels in our benchmarking results (Supplementary Fig. 4). We also performed a grid search over the relevant parameter spaces of the spot size and threshold parameters for both TrackMate and RS-FISH. Their results were shown using both global and per-image parameter tuning, where the former used a single parameter combination that achieved the highest mean F1 score, and the latter used the best parameter combination for each image. In our benchmarks, Piscis achieved a mean F1 score of 0.895, significantly outperforming all other algorithms. Although deepBlink (F1 = 0.863) outperformed both globally tuned TrackMate (F1 = 0.756) and RS-FISH (F1 = 0.665), it performed worse than per-image tuned TrackMate (F1 = 0.874), which represented the upper-performance limit for a human user who always chooses the optimal parameter combination for each image. In contrast, Piscis still surpassed this upper limit for both TrackMate and RS-FISH without needing any manual parameter tuning.

**Figure 3:**
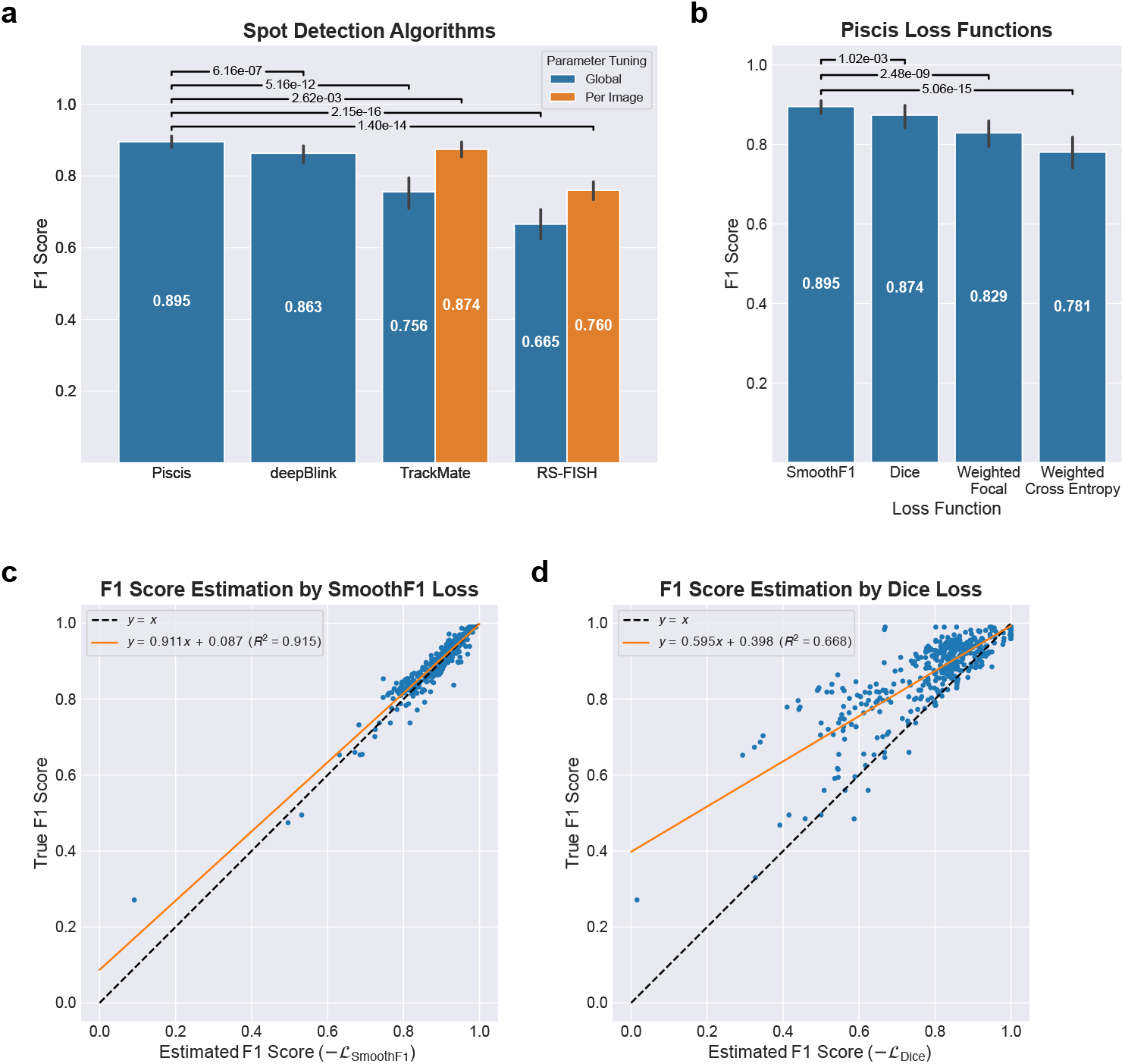
Piscis outperforms other state-of-the-art spot detection methods. **a**, Bar plot comparison between Piscis and other spot detection methods over the testing images from our combined dataset (*n* = 91). Statistical significance was determined using the one-sided Wilcoxon signed-rank test, with the hypothesis that Piscis yielded higher F1 scores. **b**, Bar plot comparison between Piscis models trained using the SmoothF1, Dice, weighted focal, and weighted cross entropy losses over the testing images from our combined dataset. Statistical significance was similarly determined using the one-sided Wilcoxon signed-rank test. **c**, Comparison between the true F1 scores and those estimated by the SmoothF1 loss for the Piscis model trained using the SmoothF1 loss. Each data point corresponds to a single image from our training dataset (*n* = 418). **d**, Comparison between the true F1 scores and those estimated by the Dice loss for the Piscis model trained using the Dice loss. Each data point again corresponds to a single image from our training dataset.

We also trained Piscis on our combined dataset using the Dice loss and two common variants of cross entropy, the weighted cross entropy loss and focal loss, to compare their performance with the SmoothF1 loss. Again, the Piscis model trained using the SmoothF1 loss, as described previously, significantly outperformed the model trained using these other common loss functions (Fig. 3b). In line with previous works that found the Dice loss to be more effective than variants of the cross entropy loss, the model trained using the Dice loss resulted in the closest performance to the SmoothF1 loss. Looking at the individual experimental and synthetic datasets included in our combined dataset, we found that training Piscis using the SmoothF1 loss yielded the highest mean F1 score for six out of nine experimental conditions and all four synthetic particle types (Supplementary Fig. 5-6).

To explain why the SmoothF1 loss resulted in better performance than the Dice loss, we plotted the F1 scores over our training set for both Piscis models against their negative loss values. Linear regression revealed that the SmoothF1 loss provided an accurate approximation of the true F1 score, as evidenced by a high *R*^2^ value of 0.915 (Fig. 3c). In contrast, the data for the Dice loss exhibited a weaker correlation, reflected in a lower *R*^2^ value of 0.668 (Fig. 3d). Furthermore, almost all points in the plot for the Dice loss lie above the *y* = *x* line, which means that it consistently underestimates the true F1 score. We believe one major cause for this underestimation is that spot pixels in the predicted classification labels can often be slightly offset from the ground truth. Even if this offset is only a single pixel, the Dice loss would classify the predicted spot pixel as a false positive. The SmoothF1 loss does not suffer from the same issue because it is computed from the pooled labels, where our deformable sum pooling operation corrects slightly offset spot pixels using the predicted displacement vectors. To demonstrate the effect of this difference, we randomly shifted spots in each image of our training set by small vectors and computed both the SmoothF1 and Dice losses. Theoretically, small offsets should still result in an F1 score of approximately 1 if the predicted displacement vectors are accurate. The SmoothF1 loss indeed has this property, whereas the Dice loss does not (Supplementary Fig. 7). The F1 scores predicted by the Dice loss rapidly decrease as the magnitude of the randomly sampled vectors increases (Supplementary Fig. 7a).

In addition to our combined dataset, we trained Piscis models on the six datasets from the deepBlink paper and compared them against the pre-trained deepBlink models. Although the differences in F1 scores were minor, Piscis (Single-molecule RNA FISH: F1 = 0.935, SunTag: F1 = 0.800, Particle: F1 = 0.950, Microtubule: F1 = 0.661, Receptor: F1 = 0.686, Vesicle: F1 = 0.739) still outperformed deepBlink (Single-molecule RNA FISH: F1 = 0.916, SunTag: F1 = 0.748, Particle: F1 = 0.943, Microtubule: F1 = 0.638, Receptor: F1 = 0.683, Vesicle: F1 = 0.729) on all datasets (Supplementary Fig. 8).

To illustrate a situation where Piscis’ superior performance was particularly advantageous, we applied all algorithms to an additional image containing a prominent autofluorescent region, which is a common feature of data collected from tissue samples (Fig. 1c). This image, featuring human melanoma cells grown in mice, was not part of our combined dataset and was analyzed after model training. As described previously, we performed a grid search to find the optimal parameter combination for both TrackMate and RS-FISH. While deepBlink tended to undercount across the image, leading to many false negatives, both TrackMate and RS-FISH yielded many false positives in the autofluorescent region. In contrast, Piscis successfully detected most true spots while avoiding the autofluorescent region (Fig. 1d). This ability to reduce false positives in noisy image regions, which normally requires extensive and tedious manual correction, demonstrates Piscis’ potential to streamline the analysis of RNA FISH-derived data.

Finally, to evaluate Piscis’ ability to scale to large volumes of images, we benchmarked its inference runtime on both CPU and GPU. We chose to perform the benchmarks on Google Colab’s T4 GPU runtime, as it is an environment freely available to anyone with a Google account. Our exact instance featured 2 virtual threads of an Intel(R) Xeon(R) CPU @ 2.00 GHz, 12.67 GB RAM, and an NVIDIA Tesla T4 GPU with 16 GB VRAM. For input images of size 256 × 256, inference took 3.34 s per image on CPU and 0.0424 s per image on GPU (Supplementary Table 2). Piscis uses DeepTile^31^, our library for large image tiling, processing, and stitching, to scale inference to arbitrarily large images. This capability is particularly crucial for applications such as modern spatial transcriptomics, where high-resolution, large-scale imaging datasets are increasingly common and require efficient processing to enable downstream analyses.

## 3 Discussion

We have here shown that Piscis is a highly capable deep learning algorithm for spot detection that significantly outperforms other state-of-the-art methods across a range of datasets representing diverse cell types and experimental conditions. It performs exceptionally well in areas of high spot density and images with high background noise, where other methods would yield many false positives.

A key to the high performance of Piscis is the development and application of a differentiable approximation of the F1 score as the loss function. The F1 score is, in many ways, the ideal loss function for many classification tasks because it naturally balances the true positive, false positive, and false negative rates. However, it is usually not differentiable, hence impossible to use for model training. Thus, our development of the SmoothF1 loss function is a significant step towards a general solution to class imbalance issues in deep learning. Indeed, the framework we provided here should allow for smooth approximations of the F1 score in many other image analysis tasks, extending the applicability of our approach to a broader range of scientific domains.

Piscis has the potential to enhance many practical aspects of spot detection. In our experience, researchers face several major problems when confronted with real-world data that create bottlenecks in the analysis pipeline. One is the challenge of uneven background intensity, which makes it impossible to choose a global threshold for spot detection methods based on Laplacian of Gaussian or Difference of Gaussian filters. Another is the false positives that appear when looking at, for example, tissue sections, which often have areas of bright autofluorescence that Laplacian of Gaussian or Difference of Gaussian methods misidentify as clusters of spots. Currently, these areas must be manually removed from the analysis, which is tedious for large imaging fields. Piscis alleviates both issues, allowing for automated analysis that enables image quantification on a much larger scale.

Another important application for spot detection is multiplex imaging. Typically, multiplex biomolecule detection with fluorescence microscopy is limited by the number of fluorescent dyes that can be discriminated via optical bandpass filters, which is usually around four or five. Nevertheless, a number of recent spatial transcriptomics techniques have emerged that circumvent these limitations to greatly enhance our ability to detect and quantify many more RNA species simultaneously, often into the hundreds or even thousands^4–6^. These methods typically work by reading out a sequence of fluorescent colors for each target RNA through multiple rounds of chemistry and imaging to obtain a molecular “barcode.” For instance, one type of RNA molecule may be labeled “Red-Green-Red-Blue,” while another may be labeled “Red-Blue-Red-Red.” Accurately reading these barcodes is a spot detection problem for which Piscis may have much practical potential. Moreover, it may prove useful to develop variants of spot detection algorithms like Piscis that can explicitly detect spots over these multiple rounds of imaging, potentially leveraging image information across successive rounds of imaging to read out barcodes more accurately than would be possible by just applying Piscis to a single round at a time.

## 4 Materials and Methods

### 4.1 Datasets

#### 4.1.1 Training and testing data

The dataset used to train Piscis and test all algorithms consists of experimental RNA FISH and synthetic data. For the experimental data, we performed either standard single-molecule RNA FISH or HCR RNA FISH and imaged cells using fluorescence widefield microscopy. We collected six datasets representing multiple cellular systems, including WM989 human melanoma cells^28,29^ grown *in vitro* and in NOD SCID mice^32^, human inducible fibroblast-like (hiF-T) cells^26,27^, Calu-3 human lung adenocarcinoma cells^30^, and primary human monocyte-derived macrophages (hMDMs). All images were acquired at 60x magnification and 1.4 numerical aperture using a Nikon Ti-E microscope. More specifics for each dataset, including the imaging channels, targeted genes, and probe fluorophores, can be found in Supplementary Table 1.

For the synthetic data, we took a subset of the datasets used to train deepBlink^20^. These included the “Particle” dataset, generated by the “Synthetic Data Generator” plugin in Fiji^33^, and the “Microtubule,” “Receptor,” and “Vesicle” datasets, created for the Particle Tracking Challenge at the 2012 International Symposium on Biomedical Imaging^34^.

#### 4.1.2 Annotation of experimental images

Experimental images were manually annotated using the custom software NimbusImage, the code for which can be found at https://github.com/arjunrajlaboratory/NimbusImage. During the annotation process, we identified spots based on their shape and intensity relative to the background while cross-referencing with the nuclei channel and adjacent z-levels when available. We also generally avoided ambiguous image regions, where a low signal-to-noise ratio made it difficult to obtain a definitive ground truth. The resulting annotations, including their *x* and *y* coordinates and fluorescence channels, were exported as .json files for further processing.

#### 4.1.3 Processing of annotated images

Several processing steps were then applied to prepare datasets for model training and testing. For the experimental data, manual annotations were improved to subpixel accuracy by Gaussian fitting. Each manual annotation was snapped to the corresponding image’s local maxima within a 3 × 3 pixel kernel. A three-dimensional Gaussian function was then fitted in a 3 × 3 pixel kernel around this local maxima using the curve_fit method from the SciPy package^35^, whereby the center of the fitted Gaussian was taken to be ground truth coordinates. Multiple annotations could be inadvertently fitted to the same spot due to accidental double annotation or high spot density. To account for these scenarios, we removed potential duplicates by replacing each group of points within a 1-pixel radius with a single point representing the centroid. After this process, images were tiled to create smaller crops of 256 × 256 pixels while keeping only those containing at least one ground truth spot. The tiled images and annotations were then randomly partitioned into training (70%), validation (15%), and testing (15%) splits to generate a single .npz file for each channel of each dataset. In the end, we obtained a total of 358 images across the six experimental datasets, as shown in Supplementary Table 1.

For the synthetic data, a subset of 60 images and their corresponding annotations were taken from each of the four deepBlink datasets, using the same split ratios as before, with 42 for training, 9 for validation, and 9 for testing. Smaller crops of 256 × 256 pixels were then extracted from the center of each image. In the end, we obtained a total of 240 images across the four synthetic datasets.

Finally, the experimental and synthetic images, along with their annotations, were compiled into our combined dataset for model training and testing.

#### 4.1.4 Additional data with autofluorescent noise

In addition to the training and testing data, we used one more dataset to evaluate the performance of all algorithms on an image with pronounced autofluorescent noise (Fig. 1c-d). For this experimental data, we performed clampFISH 2.0^9^ on WM989 cells^28^ grown in NOD SCID mice^32^ and imaged the resulting tissue section using fluorescence widefield microscopy. These images were acquired at 20x magnification and 0.75 numerical aperture using a Nikon Ti-E microscope. The specific image used in our comparison was a 512 × 512 crop from the CY3 channel, where the gene *NGFR* was labeled using probes with the fluorophore Cy3. We manually annotated this image using NimbusImage and applied the same Gaussian fitting process as previously described.

### 4.2 Piscis algorithm design

#### 4.2.1 Auxiliary representation of spots

Spots can be represented using a set of feature maps with the same spatial dimensions as the raw image, which consists of binary classification labels of each pixel as background or spot and displacement vectors pointing each pixel to the nearest true spot center (Fig. 1e). Ground truth classification labels **L** are generated by first setting pixels containing a spot center to a value of 1 and all other pixels to a value of 0. Morphological dilation can be optionally applied to expand the region occupied by each spot, mitigating class imbalance and increasing the tolerance for slight positional offsets between a model’s prediction and the ground truth. Ground truth vector displacements, with vertical components **V** and horizontal components **H**, are generated by assigning each pixel to the nearest spot center via a Voronoi tessellation and calculating their coordinate difference. A deep neural network is then used to predict these feature maps from the raw image.

#### 4.2.2 Deep neural network

The deep neural network was based on the Feature Pyramid Network^24^, a state-of-the-art architecture for object detection (Fig. 1e, Supplementary Fig. 2). In the bottom-up pathway, a modified EfficientNetV2^25^ backbone processes a one-channel input image, standardized to zero mean and unit variance, to extract four sets of feature maps **C**_1_, **C**_2_, **C**_3_, and **C**_4_ with 32, 64, 128, and 256 channels respectively. While **C**_1_ preserves the spatial dimensions of the input image, the convolutional blocks generating **C**_2_, **C**_3_, and **C**_4_ are each preceded by a downsampling operation that reduces the spatial resolution by a factor of two. This downsampling is performed via a max pooling operation instead of using a convolutional stride of two like in the original EfficientNetV2 implementation. The following stages of the network are primarily adapted from Cellpose^15^, a popular deep learning algorithm for cellular segmentation. A difference in our network architecture is the construction of a feature pyramid in the top-down pathway and the aggregation of feature maps from all spatial resolutions, in contrast to Cellpose’s utilization of only the final output layer.

Between the bottom-up and top-down pathways, an image style vector **s** is computed by a global average pooling operation on the feature maps **C**_4_ and subsequent normalization:

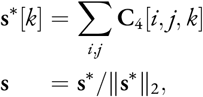

where *i, j* are the spatial indices, and *k* is the channel index. The resulting 256-dimensional unit vector **s** is a compressed representation of the input image, which is later transferred to each level in the feature pyramid.

In the top-down pathway, pyramid levels are created via a combination of upsampling and convolutional operations. First, define a 2 × 2 nearest-neighbor upsampling operation 𝒰_2×2_; a projection operation 𝒫_1×1_ combining batch normalization and a 1 × 1 convolutional layer; a convolutional operation ℱ combining batch normalization, the swish activation function, and a 3 × 3 convolutional layer; and an addition operation 𝒢of a broadcasted projection of the image style vector **s** via a dense layer 𝒫 to the output of ℱ:

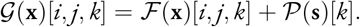

Using these operations, construct two residual blocks:

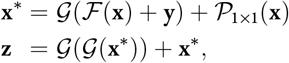

which combined form a convolutional operation ℋ with **z** = ℋ(**x, y**) (Supplementary Fig. 2b). The four pyramid levels **P**_4_, **P**_3_, **P**_2_, and **P**_1_, each with 32 channels, are then iteratively generated:

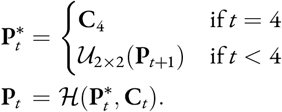

Here, setting **y** = **C**_*t*_ and adding it with ℱ(**x**) in the first residual block would create a lateral skip connection that enhances the feature maps in the top-down pathway with the equivalent resolution feature maps from the bottom-up pathway.

The top three pyramid levels **P**_4_, **P**_3_, and **P**_2_ are further upsampled to match the spatial size of the bottom level **P**_1_. First, define a combined upsampling and convolutional operation

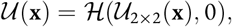

where the 0 element indicates the lack of a lateral skip connection. This operation is then applied for three iterations on **P**_4_, two iterations on **P**_3_, and one iteration on **P**_2_ to generate 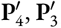, and 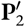 respectively:

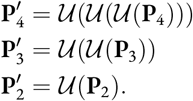

The final pyramid levels are aggregated via addition, the result of which is then projected from 32 to 3 channels, corresponding to the classification labels 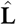 and the vertical components 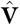 and the horizontal components 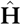 of the displacement vectors:

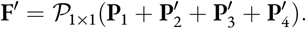

#### 4.2.3 Deformable sum pooling

A deformable sum pooling operation was applied on the predicted classification labels 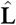 to obtain a more refined feature map we call the pooled labels 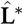, which is characterized by sharper peaks at spot centers (Fig. 1f). We implemented a discrete variant for inference that is less computationally costly and a smooth, differentiable variant for training that is compatible with gradient descent. Both variants of the operation are applied pixel-wise using a *d* × *d* pixel kernel with radius *r* = (*d* − 1)/2. During both model training and inference, we chose the radius *r* = 1, corresponding to a 3 × 3 pixel kernel, which was well-suited for the spot sizes in our dataset and minimized the computational cost. Similar to a standard convolution operation, the feature maps 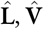, and 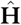 are understood to have been zero-padded on each side by *r* pixels.

For the discrete variant, define an indicator function

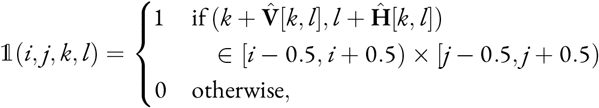

which determines whether the displaced coordinates of pixel [𝓀,*l*] falls within the bounds of pixel [*i, j*]. The pooled labels 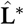 is then generated as follows:

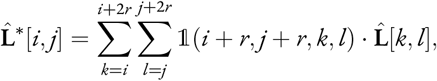

which is more sharply peaked at spot centers, enabling further post-processing steps to accurately compute their coordinates.

For the smooth variant, define a two-dimensional isotropic Gaussian function *g* with standard deviation *σ*:

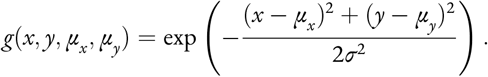

This Gaussian is used to define a soft indicator function

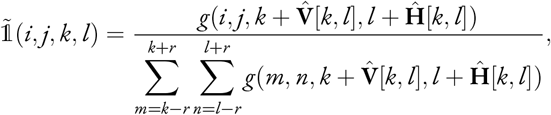

which approximates 1 by first evaluating the Gaussian, centered around the displaced coordinates of pixel [𝓀, *l*], at pixel [*i, j*]. This value is then normalized by the sum of the Gaussian over a *d* × *d* pixel kernel centered at pixel [𝓀, *l*]. Here, it is assumed that pixel [*i, j*] is contained within this kernel. To ensure that 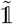 is an accurate approximation of 1, we chose *σ* = 0.5, corresponding to a narrow Gaussian with a full width at half maximum of around one pixel. The approximate pooled labels 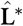 is then generated as follows:

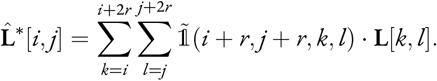

This is identical to the discrete variant except for the differences in their indicator functions.

#### 4.2.4 Post-processing of feature maps

In the post-processing step, local maxima analysis is performed on the pooled labels to identify spot center pixels with values above a fixed global threshold. An appropriate value for this threshold was determined after model training. The coordinates of these spot center pixels are then shifted by their corresponding displacement vectors to obtain predicted spot coordinates with subpixel localization accuracy (Fig. 1g).

#### 4.2.5 SmoothF1 loss function

The SmoothF1 loss function computes a differentiable approximation of the actual F1 score using the predicted pooled labels 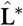 and displacement vectors 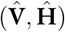 (Fig. 2b). Under the assumption that the neural network can predict pooled labels 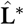 with uniformly sharp peaks at each spot, the counts of true positives (TP), false positives (FP), and false negatives (FN) are approximately proportional to the sums of label values near (*L*_TP_), distant from (*L*_FP_), and missing at (*L*_FN_) ground truth spots respectively. To obtain these values, we first compute a proximity score **S**_**D**_ for each pixel reflecting the distance between the predicted displacement vectors 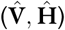 and the ground truth (**V, H**):

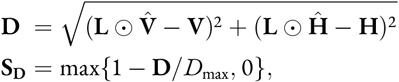

where ⊙ denotes an element-wise product and *D*_max_ is a maximum distance threshold. We chose *D*_max_ = 3 pixels, but this parameter can be adjusted to either increase or decrease the loss function’s tolerance for localization errors. It is important to note that predicted displacement vectors were “activated” only within spot pixels, specified by the ground truth classification labels **L**, to promote sparsity in the predicted feature maps 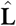 and 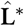. Pixels where the predicted and ground truth displacement vectors perfectly coincide will have a proximity score of 1, which decreases linearly to 0 with increasing distance until *D*_max_. The element-wise max function ensures that all pixels where **D** *> D*_max_ will have a proximity score of 0. These proximity scores are used to compute *L*_TP_, *L*_FP_, and *L*_FN_:

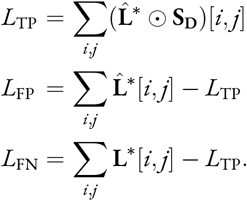

Finally, define the SmoothF1 loss function as

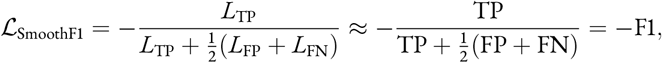

where a negative sign was introduced as loss functions by convention decrease in value as model performance improves, while the opposite is true for the F1 score.

#### 4.2.6 Masked L2 loss function

The masked L2 loss function computes the L2 norm of the difference between the predicted displacement vectors 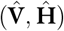 and the ground truth (**V, H**) but restricted to regions of interest defined by a binary mask. We chose the mask to be the ground truth classification labels **L**, which effectively enforces the accuracy of the predicted displacement vectors over a small region around each true spot center. The size of this region is determined by the number of morphological dilation iterations used to generate **L**. Define the masked L2 loss function as

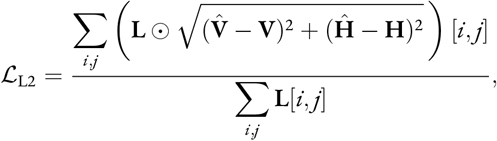

where the L2 norm of the vector difference 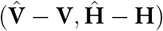 is averaged over the spot pixels in **L**.

#### 4.2.7 Composite SmoothF1-L2 loss function

For model training, define a composite SmoothF1-L2 loss function that combines the SmoothF1 and masked L2 losses:

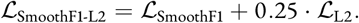

While the SmoothF1 loss alone is sufficient for training, adding the masked L2 loss as a regularization term improves training stability and model robustness. A small weighting factor of 0.25 for the ℒ_L2_ term was chosen to ensure that the ℒ_SmoothF1_ term remains the primary optimization objective. Furthermore, one morphological dilation was applied when generating the ground truth classification labels **L** to increase the loss function’s tolerance for small localization errors within a 3 × 3 region around each true spot center.

#### 4.2.8 Composite Dice-L2 loss function

To compare the SmoothF1 loss to the Dice loss used in training deepBlink, define it as

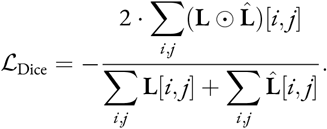

Then, define a composite Dice-L2 loss function that combines the Dice and masked L2 losses:

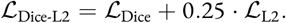

In contrast to Piscis, deepBlink adopted a grid cell strategy for spot detection, where the resolution of the network’s output feature maps was inversely proportional to the grid cell size. No morphological dilation was applied when generating the ground truth classification labels **L** to mimic the deepBlink variant using 1 × 1 pixel grid cells, which makes pixel-wise predictions from feature maps with the same spatial resolution as the raw image.

#### 4.2.9 Composite Weighted Cross Entropy-L2 loss function

To compare the SmoothF1 loss to the weighted cross entropy loss, define it as

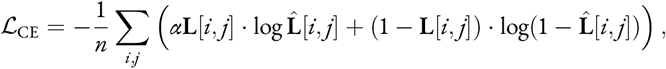

where *n* is the total number of pixels in **L** and 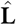 and *α* is the positive class weight. To ensure that each class contributes equally to the loss, we set

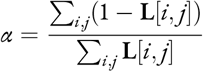

as the ratio of the negative class size to the positive class size. We expect *α* ≫ 1 in a class-imbalanced image with substantially more negative than positive pixels. Then, define a composite Weighted Cross Entropy-L2 loss function that combines the weighted cross entropy and masked L2 losses:

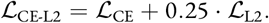

#### 4.2.10 Composite Weighted Focal-L2 loss function

To compare the SmoothF1 loss to the weighted focal loss, define it as

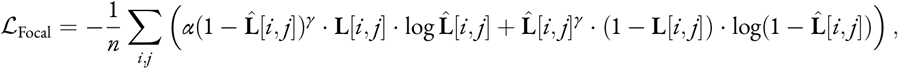

where *n* is the total number of pixels in **L** and 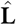, *α* is the positive class weight, and *γ* ≥ 0 is the focusing parameter. We used the same *α* as in our definition of the weighted cross entropy loss, where *α* is the ratio of the negative class size to the positive class size. We also used the default *γ* = 2 as in the focal loss paper. Then, define a composite Weighted Focal-L2 loss function that combines the weighted focal and masked L2 losses:

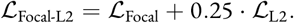

### 4.3 Training

#### 4.3.1 Data augmentation

During each training epoch, images and their corresponding ground truth annotations were randomly transformed to increase the size of the dataset artificially and prevent overfitting. First, images were standardized to zero mean and unit variance. Subsequently, each image and its annotations were randomly flipped along the horizontal and vertical axes. A random affine transformation was then applied, generated by uniformly sampling a scale from 0.75 to 1.25, a rotation angle from 0 to 2*π*, and translations in both the horizontal and vertical directions within the range − (𝓁_*s*_ − 𝓁_*f*_)*/*2 to − (𝓁_*s*_ − 𝓁_*f*_)*/*2, where 𝓁_*s*_ is the scaled image size and 𝓁_*f*_ is the model’s input size. A 𝓁_*f*_ ×𝓁_*f*_ pixel crop was then extracted from the center of the affine transformed image and annotations. Finally, each image was adjusted by a random intensity scaling factor, sampled from a log-uniform distribution supported between 0.2 and 5. This step was crucial to enhance the model’s robustness to images with skewed intensity profiles, particularly those with aberrant high-intensity pixels. The resulting augmented images and annotations containing at least one true spot were then randomly split into training batches and used to generate the ground truth classification labels ℒ and displacement vectors (**V, H**).

#### 4.3.2 Optimizer and learning rate schedule

Piscis was trained using a stochastic gradient descent optimizer with Nesterov acceleration^36,37^, a momentum of 0.9, and a weight decay of 0.0001. This optimizer was paired with a learning rate schedule, which included warmup and decay phases. During the warmup phase, accounting for the first 5% of epochs, the learning rate was linearly increased from 0 to 0.2. This rate was then held constant for the subsequent 45% of epochs. During the decay phase, accounting for the final 50% of epochs, the learning rate was halved every 5% of the epochs for a total of 10 step decays.

#### 4.3.3 Piscis models

We trained four Piscis models on our combined dataset of experimental and synthetic images using the composite SmoothF1-L2, Dice-L2, Weighted Cross Entropy-L2, and Weighted Focal-L2 loss functions. Each model was trained for 400 epochs using a batch size of four images, with every image sized at 256 × 256 pixels. Additionally, we trained six more Piscis models, each on one of the six datasets from the deepBlink paper, using the composite SmoothF1-L2 loss function. Mirroring deepBlink’s training specifications, each Piscis model was trained for 200 epochs with images sized at 512 × 512 pixels. However, unlike the deepBlink models, we used a batch size of four images instead of two to ensure stable convergence, given the increased complexity of the SmoothF1 loss function.

#### 4.3.4 deepBlink models

We retrained deepBlink on our combined dataset. Three deepBlink models were trained using 1 × 1, 2 × 2, and 4 × 4 pixel grid cells, each for 400 epochs, with all other training parameters set at their default values. Additionally, the six pre-trained models, each trained on one of the six deepBlink datasets, were downloaded and used as-is without further modifications.

### 4.4 Benchmarking

#### 4.4.1 F1 integral score

Spot detection and localization performance were evaluated using the F1 integral score^20^, referred to elsewhere as simply the F1 score. Briefly, 50 evenly-spaced numbers from 0 to 3 pixels were generated as distance thresholds. For each threshold, predicted and ground truth spots with an L2 distance below the given threshold were matched by solving a linear sum assignment problem using the Jonker–Volgenant algorithm^38^, which provided counts of true positives (TP), false positives (FP), and false negatives (FN). These counts were used to calculate the per-threshold F1 score:

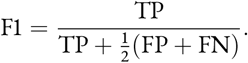

The F1 integral score was then computed by integrating per-threshold F1 scores using the trapezoidal rule and subsequently normalizing by a factor of 3 to produce a final value between a minimum of 0 (worst performance) and a maximum of 1 (best performance).

#### 4.4.2 Piscis model benchmarking

Before benchmarking the Piscis models trained using the SmoothF1 loss function, we needed to choose a global threshold for the post-processing step. To this end, the mean F1 score was computed on the validation set of each of the six deepBlink datasets using their corresponding Piscis models. A search was subsequently performed across 18 evenly-spaced threshold values from 0.5 to 9 to find the threshold that maximized the sum of the six mean F1 scores. The optimal global threshold, determined to be 1, was set as the default and used for all benchmarking and visualization purposes. With this global threshold, Piscis models trained on our combined dataset and the six deepBlink datasets were then benchmarked on their corresponding testing sets by the F1 score. Only images in the testing set containing at least one true spot were included in the final benchmarking results.

Additional Piscis models were trained using the Dice loss, weighted cross entropy loss, and weighted focal loss. A global threshold of 0.5 was set as the default for the Dice and weighted focal losses. Without a mechanism to down-weight the loss contribution of easily detected spots, the weighted cross entropy loss generally yields much higher classification label values and, in practice, requires a higher threshold to avoid false positives. Following the implementation of Polaris, we set a global threshold of 0.95 as the default for the weighted cross entropy loss.

The SmoothF1 and Dice losses were further compared to explain the differences in their performance. Piscis models trained with each loss function were applied to the training set of our combined dataset, the result of which was used to compute the true F1 scores and the corresponding loss values. For each Piscis model, negative loss values were taken as the predicted F1 scores and plotted against the true F1 scores. Linear regression was performed on the data in these two scatter plots to determine the ability of the SmoothF1 and Dice losses to approximate the true F1 score.

The tolerance of each loss function to minor mistakes, particularly for small offsets of spot pixels between the predicted and ground truth classification labels, was also tested by first shifting spots in each image of our training set by random vectors. These vectors were sampled from a two-dimensional isotropic Gaussian distribution with covariance Σ = *σ*^2^*I*. The randomly shifted spots were then used to generate classification labels with spot pixels slightly offset from the ground truth, meant to mimic a hypothetical model output. Under the assumption that a hypothetical model can still accurately predict the displacement vectors, the SmoothF1 and Dice losses were recomputed using the newly generated classification labels. This procedure was repeated for 101 evenly-spaced values of *σ* from 0 to 1.

#### 4.4.3 deepBlink model benchmarking

deepBlink models were evaluated in the same way as the Piscis models. It is important to note that our results for the six pre-trained deepBlink models deviate slightly from the original deepBlink benchmarks, which included all images in the testing set, even those containing no spots. Among the three deepBlink models trained on our combined dataset using different grid cell sizes, the model with 2 × 2 pixel grid cells significantly outperformed the other two and was selected for all comparisons with other algorithms (Supplementary Fig. 4).

#### 4.4.4 TrackMate and RS-FISH benchmarking

TrackMate and RS-FISH were each benchmarked on the testing set of our combined dataset using 100 different parameter combinations. For TrackMate, a grid search was performed across 5 evenly-spaced radius values from 1 to 3 pixels and 20 evenly-spaced threshold values from 0.02 to 0.4. Likewise, for RS-FISH, a grid search was performed across 5 evenly-spaced sigma values from 1 to 3 and 20 evenly-spaced threshold values from 0.002 to 0.04.

#### 4.4.5 Statistical analysis

The statistical significance of differences in the benchmarking results between algorithms was determined by the one-sided Wilcoxon signed-rank test, with the hypothesis that Piscis yielded higher F1 scores.

### 4.5 Software implementation

The Piscis code library was written in Python 3^39^ and leverages numerous open-source Python packages, including DeepTile^31^, Flax^40^, JAX^41^, Numba^42^, NumPy^43^, OpenCV^44^, Optax^45^, pandas^46,47^, scikit-image^48^, SciPy^35^, and Xarray^49,50^. The custom deformable pooling operation and loss functions were implemented using JAX, a high-performance numerical computing library from Google Research. The model architecture was implemented using Flax, a neural network library for JAX. The stochastic gradient descent optimizer used for model training was implemented using Optax, an optimization library for JAX. During model inference, DeepTile facilitates the processing of large image inputs by splitting them into smaller tiles. These tiles are processed in parallel, and their outputs are subsequently stitched back together, allowing Piscis to scale effectively to images of arbitrary sizes.

## 5 Data availability

Our combined dataset for model training and testing can be found at https://huggingface.co/datasets/wniu/Piscis/tree/main/20230905.

## 6 Code availability

The Piscis code library can be found at the GitHub repository https://github.com/zjniu/Piscis. Within the same repository, all code for generating the figures in this manuscript can be found at https://github.com/zjniu/Piscis/tree/main/paper. The accompanying documentation for the code library can be found at https://piscis.netlify.app. All pre-trained Piscis models can be found at the HuggingFace repository https://huggingface.co/wniu/Piscis.

## 7 Author Contributions

Z.N. and A.R. conceived the project. Z.N. designed and implemented computational methods, trained deep learning models, and performed data analysis. A.O., J.L., S.R., N.J., I.D., and Y.G. acquired experimental data for model training and testing. Z.N., A.O., J.L., S.R., and N.J. manually annotated images. Z.N. and A.R. wrote the manuscript.

## 9 Acknowledgements

The authors thank members of the Arjun Raj Lab, particularly Laila Norford, Gianna Busch, and Luciann Cuenca, for insightful discussions and feedback related to this work. The authors thank Gianna Busch, Ryan Boe, Catherine Triandafillou, Yael Heyman, and Alexandre Tourneux for helping with image annotation. The authors thank David Van Valen for insightful feedback. The authors thank Emily Cento, Zhilin Chen, Max A. Eldabbas, and Emileigh Maddox of the Human Immunology Core and the Division of Transfusion Medicine and Therapeutic Pathology at the Perelman School of Medicine at the University of Pennsylvania for providing de-identified monocytes that were purified from healthy donor apheresis using StemCell RosetteSep™ kits. The HIC is supported in part by NIH P30 AI045008 and P30 CA016520. HIC RRID: SCR_022380.

Z.N. acknowledges support from the Roy and Diana Vagelos Scholars Program in the Molecular Life Sciences, Roy and Diana Vagelos Science Challenge Award, Barry M. Goldwater Scholarship, Fannie and John Hertz Foundation Fellowship, Paul & Daisy Soros Fellowship for New Americans, and Department of Energy Computational Science Graduate Fellowship. Y.G. acknowledges support from the Burroughs Wellcome Fund Career Awards at the Scientific Interface. A.R. acknowledges support from NIH 4DN U01DK127405.

