## Supplementary Figures for "Piscis: a novel loss estimator of the F1 score enables accurate spot detection in fluorescence microscopy images via deep learning"

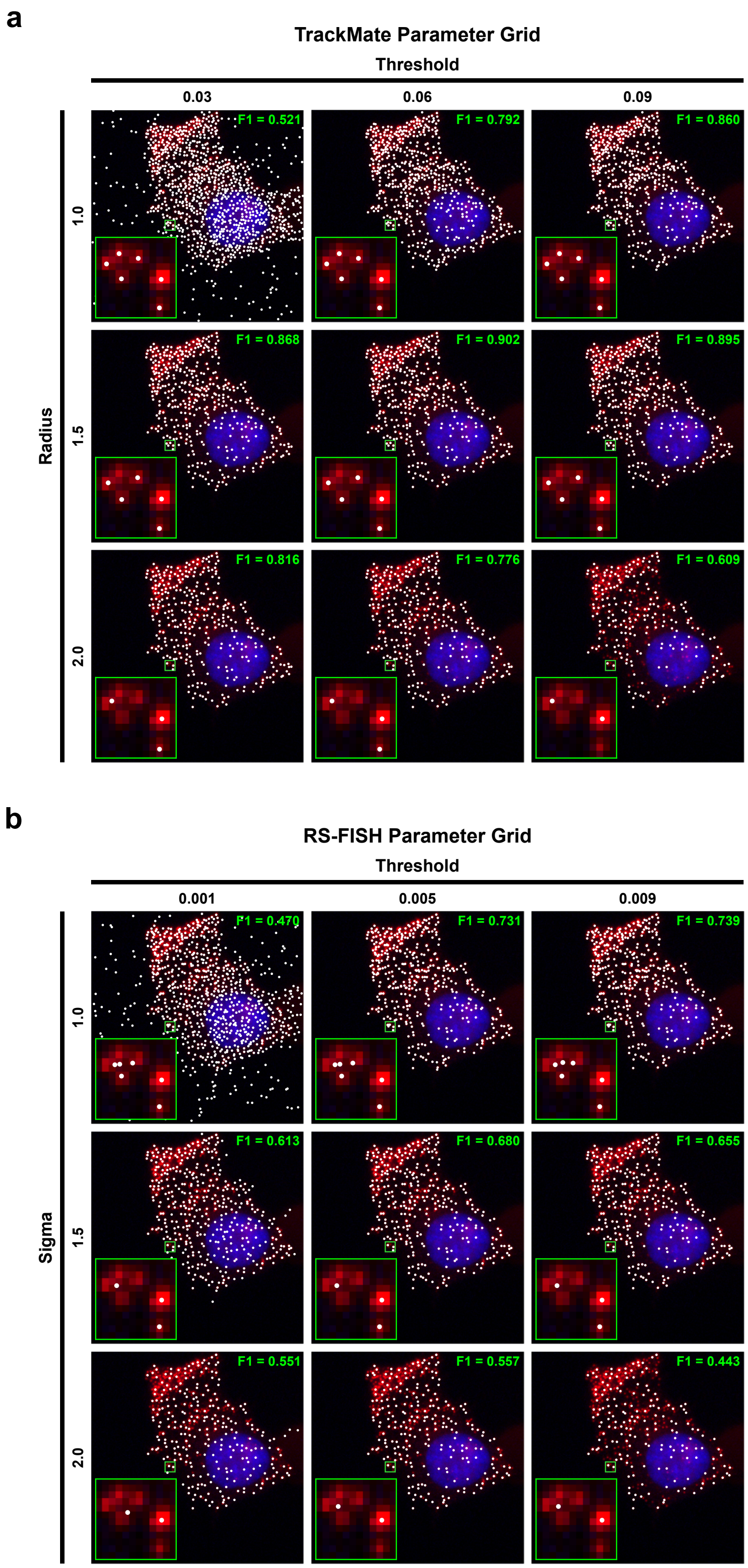


**Supplementary Figure 1**: **Traditional computational methods for spot detection are highly sensitive to parameter tuning. a,** TrackMate with different combinations of its radius and threshold parameters applied to the image from Fig. 1a. **b,** RS-FISH with different combinations of its sigma and threshold parameters applied to the image from Fig. 1a.

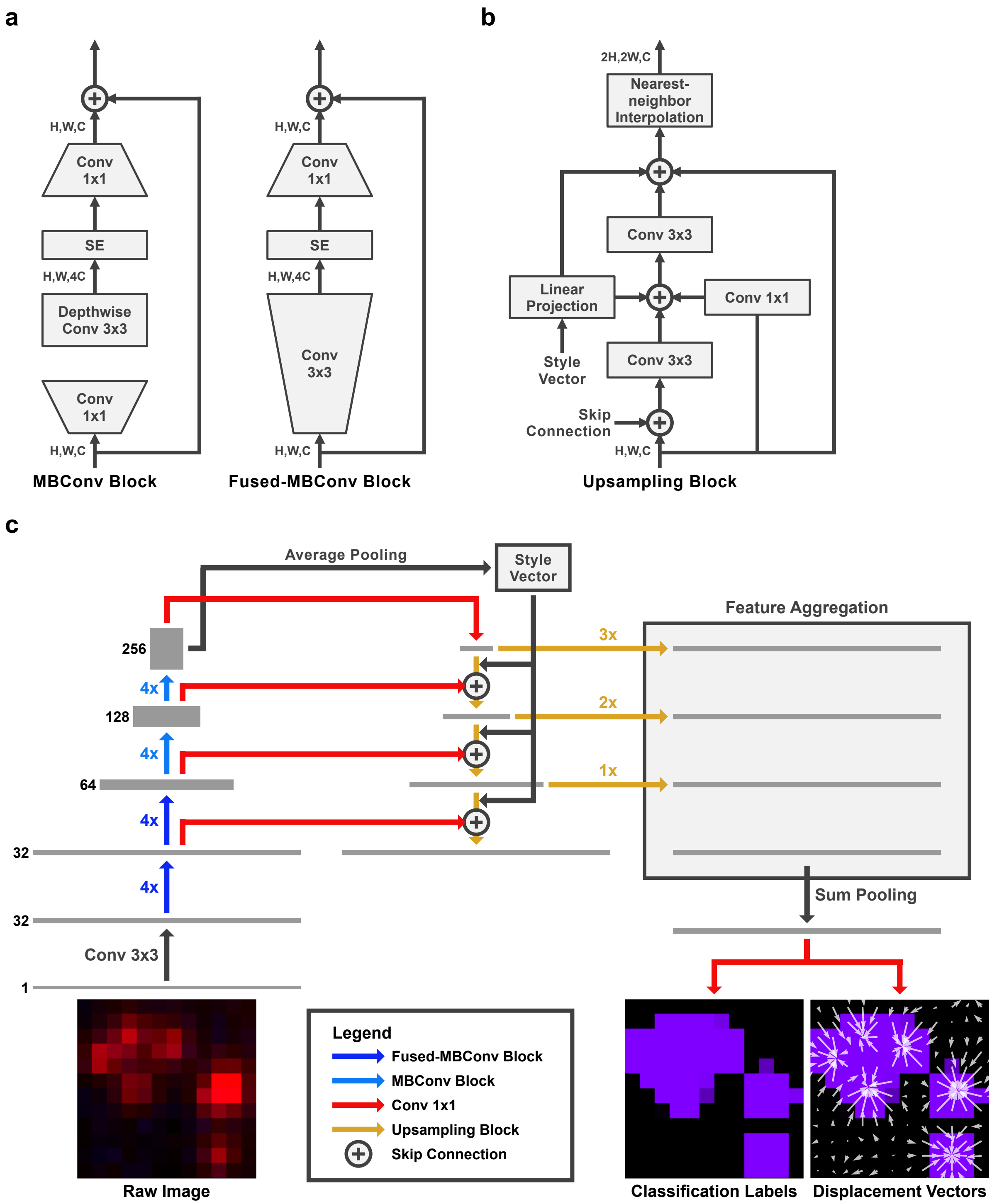


**Supplementary Figure 2: Piscis uses a Feature Pyramid Network with an EfficientNetV2 backbone to predict classification labels and displacement vectors from a raw image. a,** Building blocks of the EfficientNetV2 backbone, adapted from [25]. "SE" corresponds to Squeeze-and-Excitation blocks, which are known to strengthen the network's representational power[51]. **b,** Upsampling block combining convolution and nearest-neighbor interpolation operations. **c,** The bottom-up pathway consists of an EfficientNetV2 backbone that downsamples the raw image via repeated MBConv and Fused-MBConv blocks while progressively generating more features. Average pooling of the top-most layer generates the style vector to be used by the Upsampling block, as shown in **b**. The top-down pathway consists of Upsampling blocks and lateral skip connections via $1 \times 1$ convolutions from the equivalent-resolution feature maps in the bottom-up pathway. Lower-resolution feature maps in the top-down pathway are further upsampled to match the resolution of the raw image, and the resulting feature maps from each level in the feature pyramid are aggregated via sum pooling. An additional $1 \times 1$ convolution then generates predictions for the classification labels and displacement vectors.

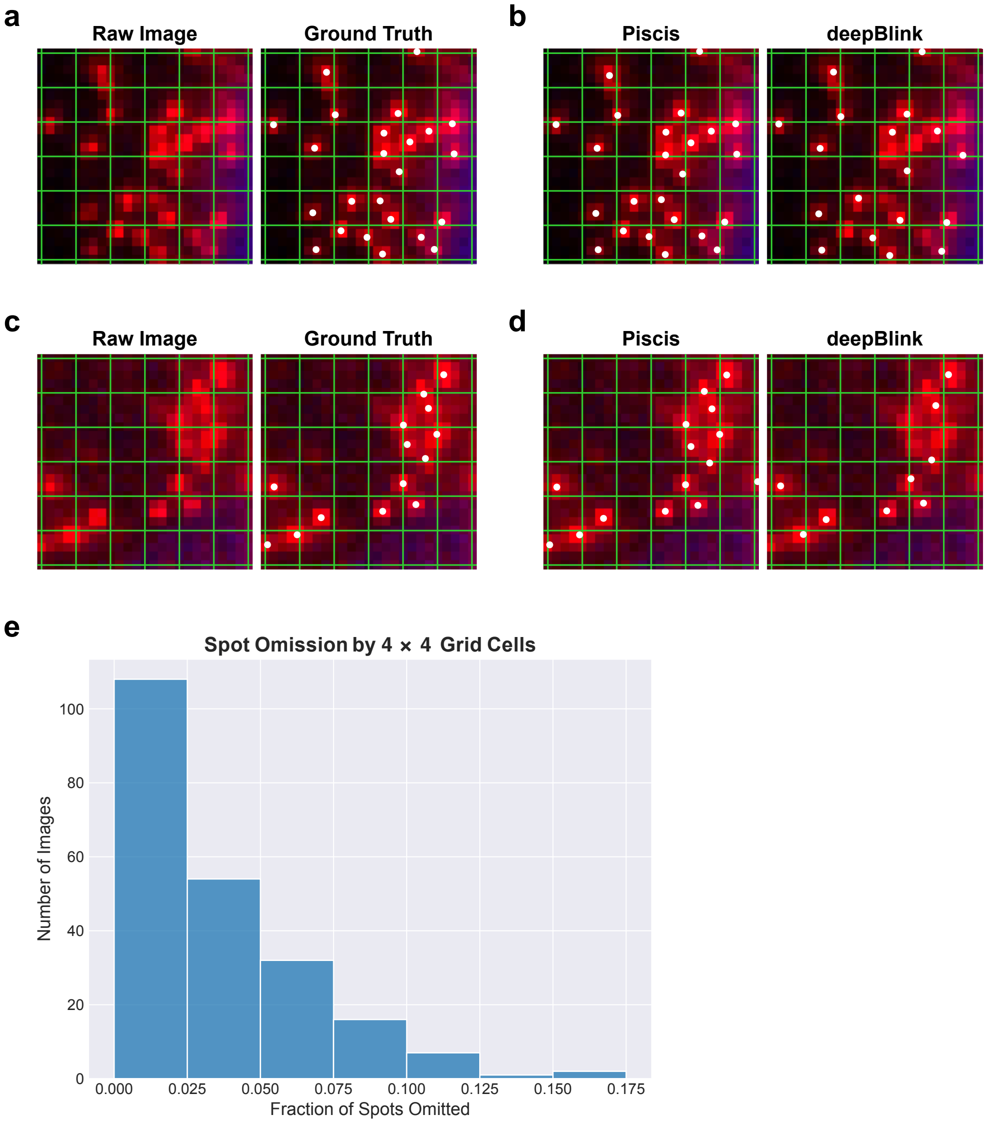


**Supplementary Figure 3: deepBlink's grid cell strategy leads to the omission of spots in regions of high spot density. a,** A single-molecule RNA FISH image of human fibroblast cells grown *in vitro* with DAPI-stained nuclei (blue), spots of single mRNA molecules for the gene *UBC* (red), and the corresponding manual ground truth annotations (white). A grid (green) with $4 \times 4$ cells is drawn over the images. **b,** Comparison between Piscis and deepBlink ($4 \times 4$ grid cells) applied to the image from **a**. **c,** A single-molecule RNA FISH image of human fibroblast cells grown *in vitro* with DAPI-stained nuclei (blue), spots of single mRNA molecules for the gene *SPP1* (red), and the corresponding manual ground truth annotations (white). A grid (green) with $4 \times 4$ cells is again drawn over the images. **d,** Comparison between Piscis and deepBlink ($4 \times 4$ grid cells) applied to the image from **c**. **e,** Histogram showing the distribution of fraction of spots omitted by $4 \times 4$ grid cells for the 220 experimental images in our dataset containing at least ten spots.

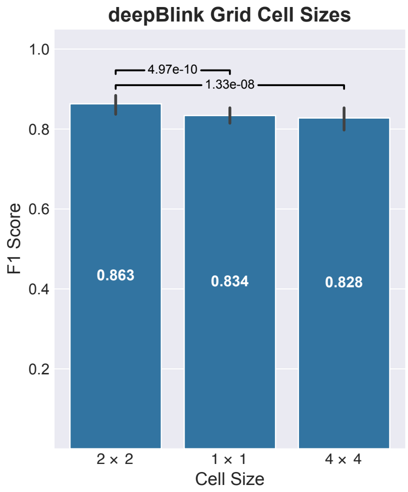


**Supplementary Figure 4**: **deepBlink's performance is dependent on its grid cell size.** Bar plot comparison between deepBlink models using different grid cell sizes over the testing images from our combined dataset ($n = 91$). Statistical significance was determined using the one-sided Wilcoxon signed-rank test, with the hypothesis that the deepBlink model using $2 \times 2$ grid cells yielded higher F1 scores.

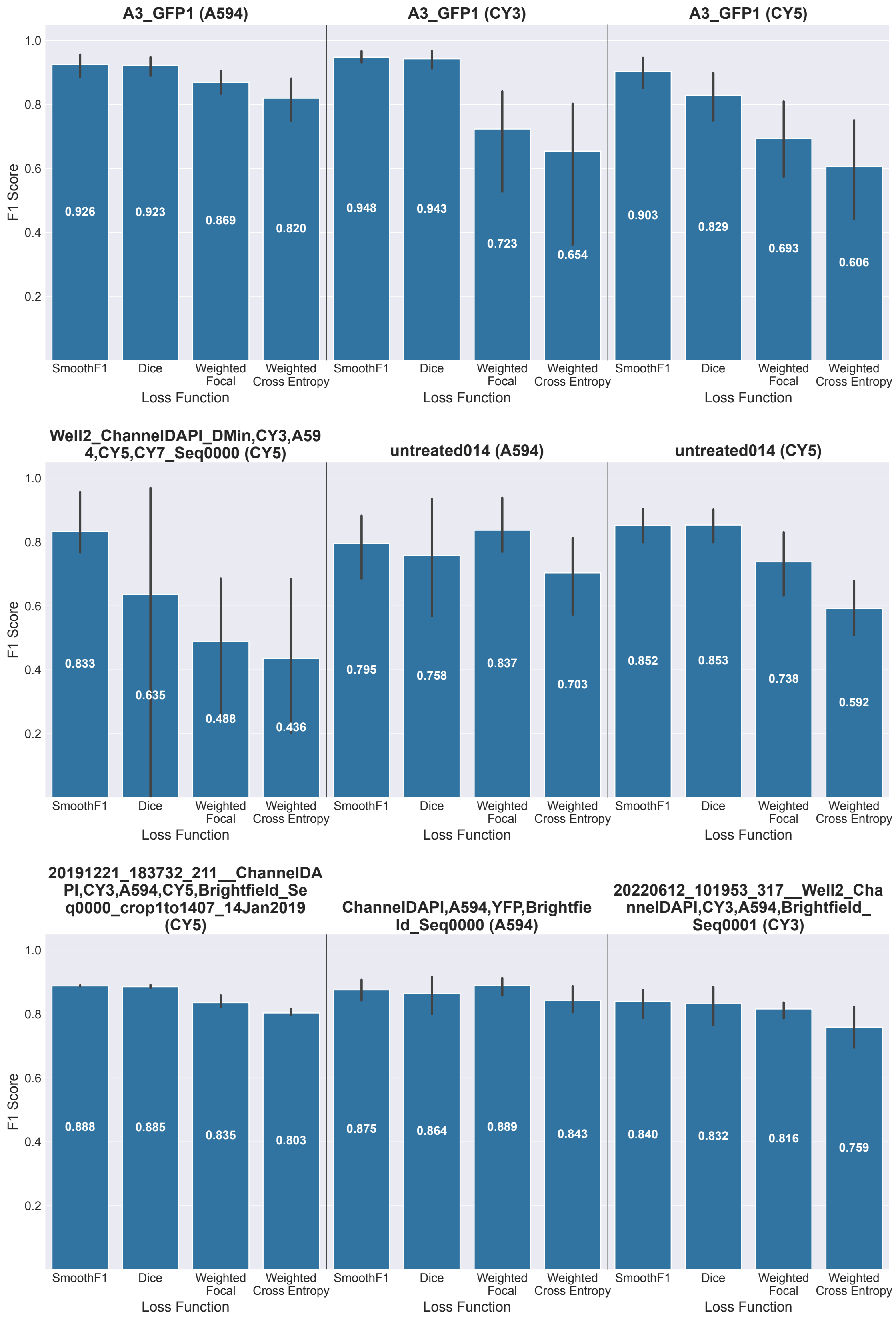


**Supplementary Figure 5: The SmoothF1 loss outperforms other loss functions across most experimental conditions in our combined dataset.** Bar plot comparison between Piscis models trained using the SmoothF1, Dice, weighted focal, and weighted cross entropy losses over the testing images across experimental conditions in our combined dataset (first row: $n = 11, 4, 6$; second row: $n = 3, 4, 14$; third row: $n = 3, 5, 5$).

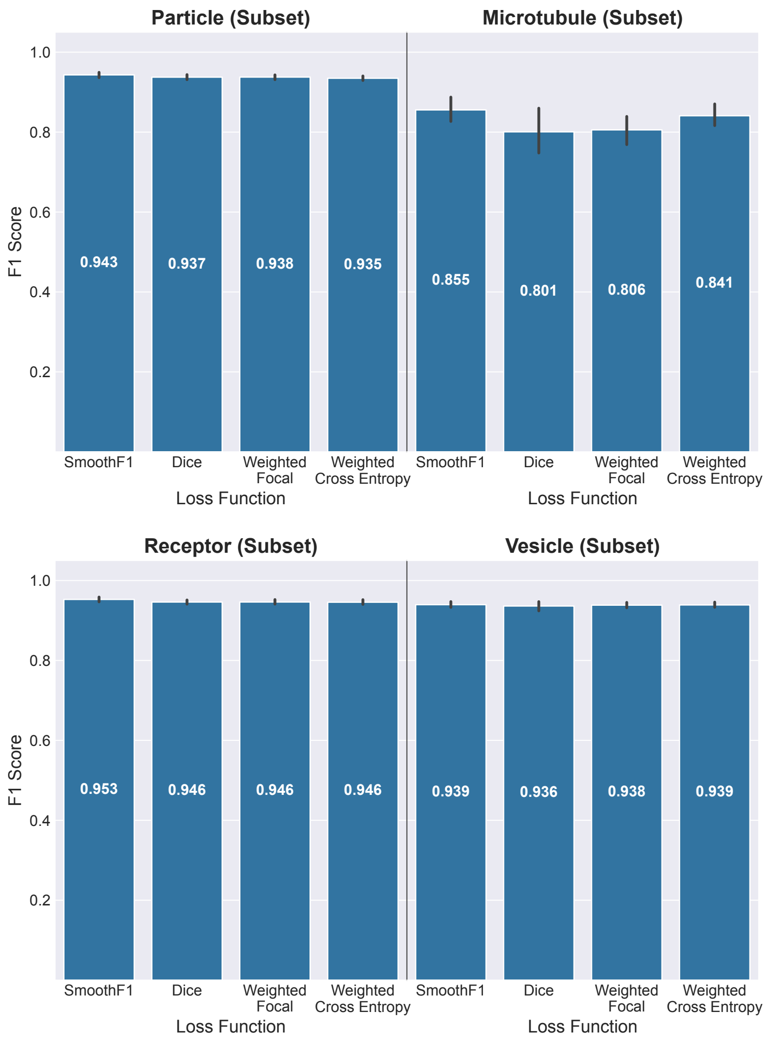


**Supplementary Figure 6: The SmoothF1 loss outperforms other loss functions across all synthetic particle types in our combined datasets.** Bar plot comparison between Piscis models trained using the SmoothF1, Dice, weighted focal, and weighted cross entropy losses over the testing images across synthetic particle types in our combined dataset ($n = 9$ for each particle type).

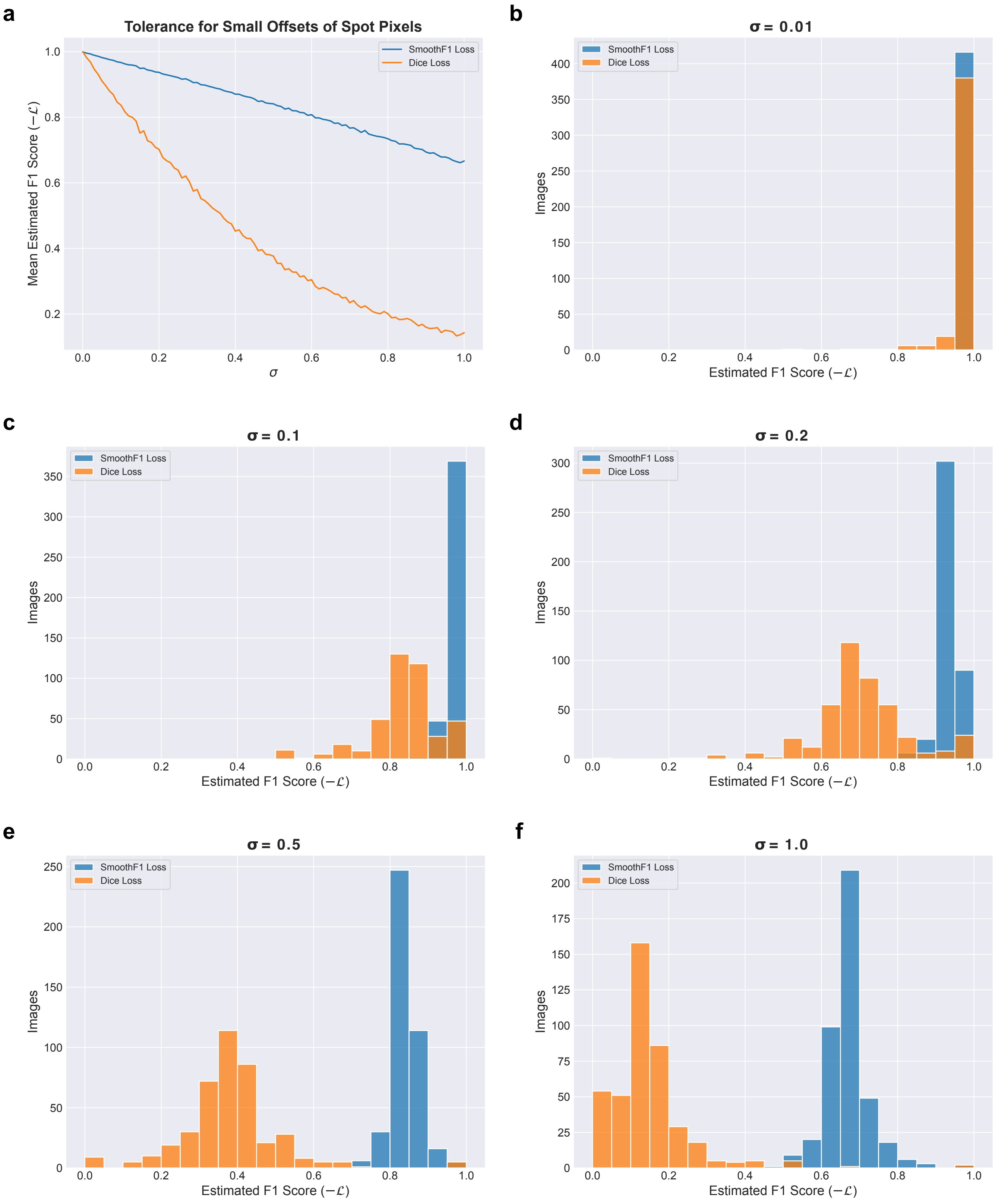


**Supplementary Figure 7: The SmoothF1 loss has more tolerance than the Dice loss for small offsets of spot pixels.** **a,** Plot of the mean F1 score estimated by the SmoothF1 and Dice losses for the training images from our combined dataset ($n = 418$) as the value of $\sigma$ for a two-dimensional isotropic Gaussian distribution with covariance $\Sigma = \sigma^2 I$, from which spot offset vectors are sampled, varies between 0 to 1. Histograms show the distribution of F1 scores for our training images when **b,** $\sigma = 0.01$; **c,** $\sigma = 0.1$; **d,** $\sigma = 0.2$; **e,** $\sigma = 0.5$; and **f,** $\sigma = 1.0$.

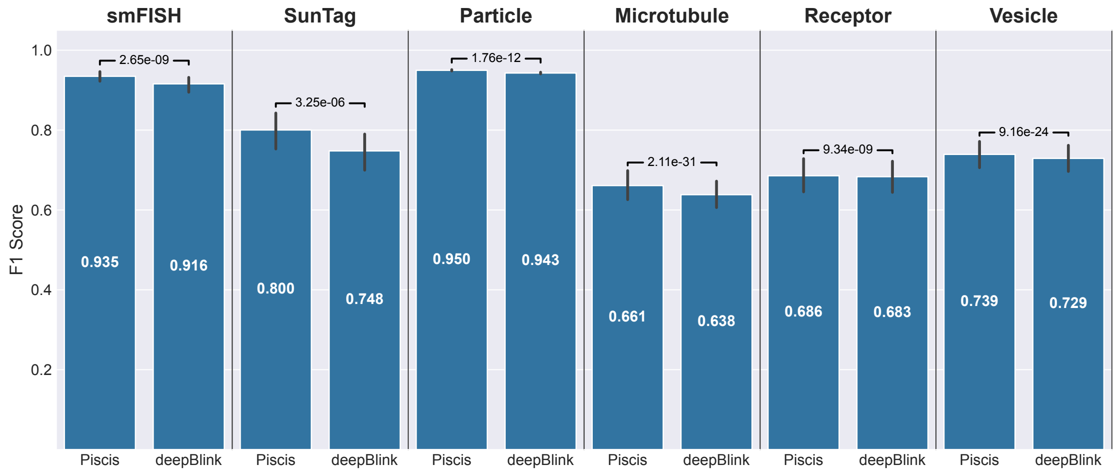


**Supplementary Figure 8**: **Piscis outperforms deepBlink on all six datasets from the deepBlink paper.** Bar plot comparison between Piscis and deepBlink over the testing images from the six deepBlink datasets: "Single-molecule RNA FISH" ($n = 129$), "SunTag" ($n = 105$) "Particle" ($n = 64$), "Microtubule" ($n = 240$), "Receptor" ($n = 240$), and "Vesicle" ($n = 240$). Statistical significance was determined using the one-sided Wilcoxon signed-rank test, with the hypothesis that Piscis yielded higher F1 scores.
