## Supplementary Tables 1-2 for "Piscis: a novel loss estimator of the F1 score enables accurate spot detection in fluorescence microscopy images via deep learning"

| Dataset Name | Method | Cell Type | Channel | Gene | Fluorophore | Images |
|---|---|---|---|---|---|---|
| 20191221_183732_211__ChannelDAPI,CY3,A594,CY5,Brightfield_Seq0000_crop1to1407_14Jan2019 | Single-molecule RNA FISH | WM989 cells (in NOD SCID mice) | CY5 | *UBC* | Atto 647N | 16 |
| 20220612_101953_317__Well2_ChannelDAPI,CY3,A594,Brightfield_Seq0001 | Single-molecule RNA FISH | hMDMs (*in vitro*) | CY3 | *UBC* | Cy3 | 32 |
| A3_GFP1 | Single-molecule RNA FISH | hiF-T cells (*in vitro*) | A594 | *UBC* | Alexa Fluor 594 | 69 |
| | | | CY3 | *MKI67* | Quasar 570 | 27 |
| | | | CY5 | *SPP1* | Quasar 670 | 40 |
| ChannelDAPI,A594,YFP,Brightfield_Seq0000 | HCR RNA FISH | Calu-3 cells (*in vitro*) | A594 | *MMP7* | Alexa Fluor 594 | 32 |
| Well2_ChannelDAPI_DMin,CY3,A594,CY5,CY7_Seq0000 | Single-molecule RNA FISH | WM989 cells (*in vitro*) | CY5 | *FKBP5* | Atto 647N | 21 |
| untreated014 | Single-molecule RNA FISH | WM989 cells (*in vitro*) | A594 | *TGM2* | Alexa Fluor 594 | 27 |
| | | | CY5 | *FKBP5* | Atto 647N | 94 |

**Supplementary Table 1**: **Experimental RNA FISH datasets included in our combined dataset.** Information provided for each dataset consists of the experimental method, cell type, imaging channels, targeted genes, probe fluorophores, and the number of resulting images after processing.

| | **CPU** | **GPU** |
|---|---|---|
| 100-image Batch | 5min 33s $\pm$ 3.64 s | 4.24 s $\pm$ 0.357 s |
| Per-image | 3.34 s | 0.0424 s |

**Supplementary Table 2**: **Piscis runtime benchmarking.** A random batch of 100 images of size $256 \times 256$ was chosen from our combined dataset for runtime benchmarking. All benchmarks were performed on Google Colab's free-tier T4 GPU runtime, which featured 2 virtual threads of an Intel(R) Xeon(R) CPU @ 2.00 GHz, 12.67 GB RAM, and an NVIDIA Tesla T4 GPU with 16 GB VRAM. Per-image runtimes were obtained by dividing the 100-image batch runtimes by 100.
